# High-integrity nanoemulsions formulation of resiquimod (R848) enhances stability and delivery for triple negative breast cancer immunotherapy

**DOI:** 10.1101/2025.04.04.647265

**Authors:** Ahmed Salem, Sara Attia, Samah El-Ghlban, A.S. Montaser, Mohamed F. Abdelhameed, Maged W. Helmy, Mohamed F. Attia

## Abstract

Triple-negative breast cancer (TNBC) poses significant clinical challenges due to its high heterogeneity, with multiple subtypes exhibiting distinct molecular characteristics and treatment responses. The development of resistance to chemotherapy and targeted therapies remains a major obstacle, and identifying reliable biomarkers to predict therapeutic response continues to be challenging. This study aims to enhance the delivery of the immunostimulant Toll-like receptor 7/8 (TLR7/8) agonist resiquimod (R848) using a safe and highly integrated nanoemulsions (NEs) formulation, providing effective and reliable immunotherapy. A series of NEs were prepared and optimized with and without the reactive lipophilic compound ricinoleic acid. Neutral and negatively charged NE formulations encapsulating R848 were compared. The physicochemical properties and in vitro delivery of resiquimod into RAW 264.7 macrophages and 4T1 TNBC cell line models were studied. Both R848-loaded NE formulations exhibited prolonged shelf-life stability with minimal protein binding. Incorporating small portions of ricinoleic acid into the formulation (negatively charged NEs) slowed drug release and improved physical properties and overall delivery compared to ricinoleic-free formulations, likely due to its interaction with the drug. Cytotoxicity and cellular uptake studies were conducted on both NE models, showing localization in the macrophage cell membrane and 4T1 cell cytoplasm. Molecular profiling in 4T1 cells revealed R848-NEs modulated key biomarkers (TLR4/7, Cyclin D1, NF-κB) while potently inducing autophagy (evidenced by LC3II/p62/Beclin-1 alterations) and PD-L1 upregulation. These dual effects—autophagy-mediated tumor suppression and immune checkpoint modulation—suggest therapeutic synergy between R848-NEs and anti-PD-L1 antibodies, presenting a promising combinatorial strategy for TNBC treatment.

**Graphical Abstract:** 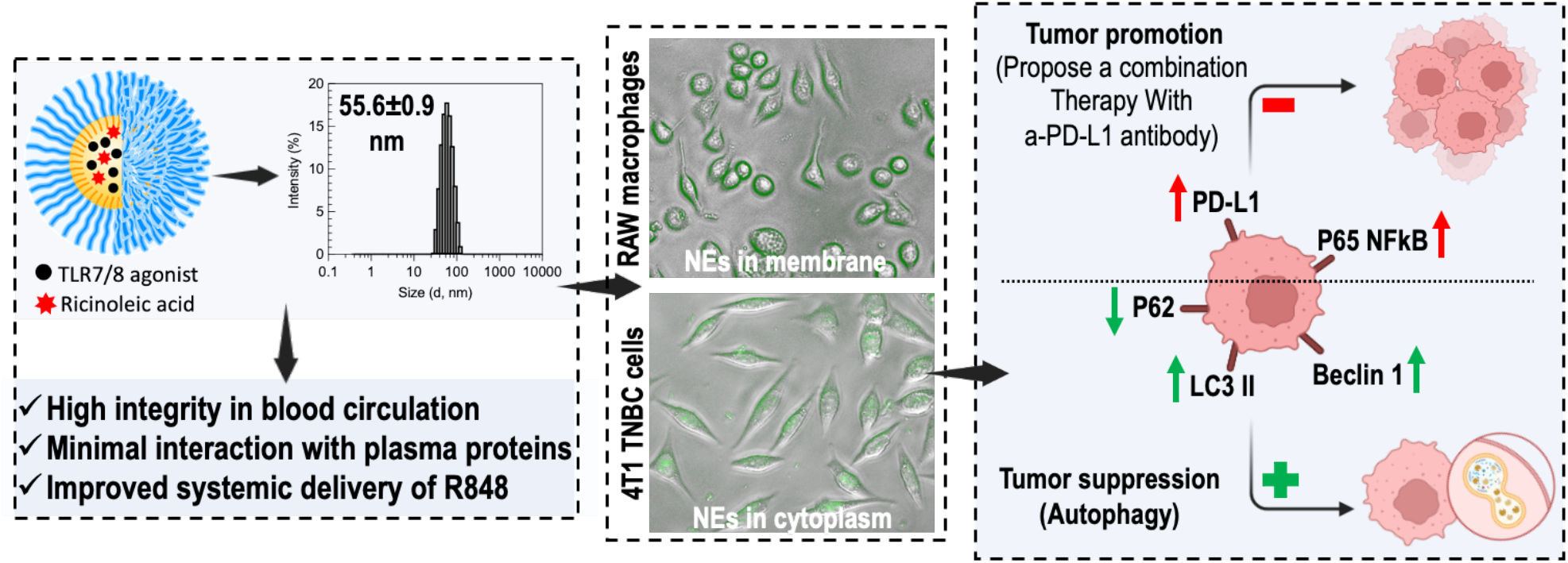

## Introduction

Triple-negative breast cancer (TNBC) is an aggressive subtype lacking estrogen (ER), progesterone (PR), and human epidermal growth factor receptor 2 (HER2) receptors, making it unresponsive to hormonal or HER2-targeted therapies [1-4]. Standard treatment is chemotherapy, but prognosis remains poor due to high recurrence and metastasis rates [5, 6]. Immunotherapy, including targeting Toll-like receptors (TLRs), offers new treatment avenues [7, 8]. TLRs are pattern recognition receptors (PRRs), a class of proteins that play a crucial role in the innate immune system. Targeting these molecules helps trigger an immune response using immunostimulant drugs [9, 10]. Resiquimod (R848) is a synthetic compound that acts as an agonist for Toll-like receptors 7 and 8 (TLR7/8) and shows promise due to its potent immunomodulatory effects in the immune tumor microenvironment (TME). It activates immune cells through the TLR7/8 MyD88-dependent signaling pathway, stimulating pro-inflammatory cytokine production and enhancing dendritic cells (DCs) and macrophages. This activation polarizes the immune response towards a Th1 phenotype, characterized by the production of interferon-gamma (IFN-γ) and other pro-inflammatory cytokines [11-13]. This reprograms TME to recruit and activate cytotoxic T lymphocytes (CTLs) and natural killer (NK) cells [9, 14, 15]. Additionally, resiquimod induces apoptosis in cancer cells through TLR7/8 signaling pathways, resulting in programmed cell death and a powerful cancer therapy agent [9, 16].

As an immune adjuvant, resiquimod enhances the body’s response to antigens, crucial for targeting and destroying tumor cells [10, 17]. This effect can improve cancer vaccine efficacy and other immunotherapeutic strategies. Resiquimod enhances antitumor immunity by activating immune cells, promoting cytokine and chemokine production, and serving as a vaccine adjuvant. Preclinical studies show resiquimod’s potential in various cancer models [18]. For instance, in melanoma, it enhances immune checkpoint inhibitors (ICIs) like anti-PD-1 therapy, improving immune response and resulting in prolonged survival and better tumor control compared to monotherapy [19-21]. Overall, resiquimod’s ability to modulate the immune system and reprogram the TME, along with its direct cytotoxic effects, underscores its potential as a tumor-modifying agent. These properties make it a promising candidate for combination therapies to enhance existing cancer treatments and overcome immunosuppressive TME challenges.

The clinical application of resiquimod faces several challenges, including poor water solubility, immunogenicity, and significant side effects such as local inflammation and systemic immune activation, which can minimize its therapeutic window [22]. Clinical trials have also shown variable efficacy, with some studies reporting limited antitumor activity [23, 24]. These challenges highlight the need for enhanced delivery methods to maximize its therapeutic potential. Nanoemulsion (NE)-based delivery systems can address these limitations by improving its systemic delivery, pharmacokinetics, bioavailability, and therapeutic efficacy. Encapsulating R848 in NEs facilitates sustained release, reduces clearance, increases half-life, and enhances antitumor efficacy, offering a comprehensive strategy for treating TNBC by reprogramming TME and inducing apoptosis in cancer cells.

This study focuses on designing, optimizing, and characterizing R848-loaded nanoemulsions (R848-NEs) for treating triple-negative breast cancer. It was found that incorporating a small amount of reactive lipophilic compound, ricinoleic acid into the NEs formulation improved its physical properties. We assessed stability, storage, plasma protein binding, and drug release kinetics for both neutral and negatively charged R848-NEs. Ricinoleic acid slowed drug release and enhanced in vitro therapeutic activity in 4T1 TNBC cell lines. While TLR7/8 agonists are known to polarize tumor-associated macrophages into the M1 phenotype, remodeling TME and immune response, our study focused on 4T1 TNBC cells to evaluate autophagy and direct tumor cell killing. The results showed autophagy induction in cancer cells and increased PD-L1 expression, suggesting that combining immune checkpoint inhibitors (i.e. a-PD-L1 or a-PD-1 antibodies) with R848-NEs could boost antitumor immunity and therapeutic response.

## Experimental section

### Chemicals and reagents

DL-α-tocopherol acetate (Vitamin E acetate, VEA), non-ionic amphiphilic PEGylated surfactant Solutol^®^ HS15 (polyethylene glycol (15)-hydroxy stearate), ricinoleic acid, and TLR7/8 agonist resiquimod were sourced from Sigma Aldrich (St. Louis, MO). Thermo Fisher Scientific supplied Hoechst 33258 solution, the lipophilic tracer DiIC18(3) (1,1’-dioctadecyl-3,3,3’,3’-tetramethylindocarbocyanine perchlorate), and cell culture reagents, including phosphate-buffered saline (PBS), Dulbecco’s Modified Eagle Medium (DMEM), fetal bovine serum (FBS), and penicillin-streptomycin solution (10,000 U/mL penicillin, 10 mg/mL streptomycin). Additional materials from Thermo Fisher Scientific comprised normal saline (0.9% NaCl), 0.22 µm syringe filters, Vybrant™ DiO Cell-Labeling Solution, Cell Counting Kit-8 (CCK-8), and 3-(4,5-dimethylthiazol-2-yl)-2,5-diphenyltetrazolium bromide (MTT). Cell lines—RAW 264.7 macrophages, 4T1 murine breast cancer cells, and Nile red-stained 4T1 (NR-4T1) cells—were obtained from the American Type Culture Collection (ATCC). Protein expression levels were quantified using commercially available ELISA kits according to manufacturers’ protocols. TLR4 levels were determined using the Mouse TLR4 ELISA Kit (Elabscience^®^, Cat. No. E-EL-M2417), while TLR7 was measured with the Mouse Toll-like Receptor 7 ELISA Kit (MyBioSource, Cat. No. MBS2533523). Cell cycle progression was assessed through Cyclin D1 quantification (Novus Biologicals, Cat. No. NBP2-75101), and NF-κB signaling activity was evaluated using the Mouse NFκB ELISA Kit (Wuhan Fine Biotech, Cat. No. EM1230). Immune checkpoint marker PD-L1 (CD274) and apoptotic marker Caspase-3 were analyzed with kits from Cusabio Biotech (Cat. Nos. CSB-EL004911MO and CSB-E08858m, respectively). Autophagy-related proteins were measured using the SQSTM1/p62 ELISA Kit (Thermo Fisher Scientific, Cat. No. A246887), LC3II ELISA Kit (MyBioSource, Cat. No. MBS3802393), and Rat Beclin 1 ELISA Kit (MyBioSource, Cat. No. MBS2706719) to comprehensively evaluate autophagic flux.

## Methods

### Formulation of R848 drug loaded-nanoemulsions (R848-NEs)

The nanoemulsions (NEs) were prepared using a low-energy spontaneous emulsification method [25-28]. The oil phase, comprising VEA, Solutol^®^ HS15 PEGylated surfactant, with or without ricinoleic acid and R848 drug, was heated to 50–60 °C for 5 minutes and vortexed until fully dissolved and clear. The aqueous phase, normal saline (0.9% NaCl), was immediately added and vortexed for 2–3 minutes, forming a colloidal nanosuspension of NEs and R848-NEs. Formulations were designed with varying component ratios, as shown in **Figure 1C**. The samples were filtered through a 0.22 *µ*m syringe filter for sterilization and to remove any free molecules. They were then stored at either room temperature (RT) or 4 °C for further analysis.

**Figure 1.**
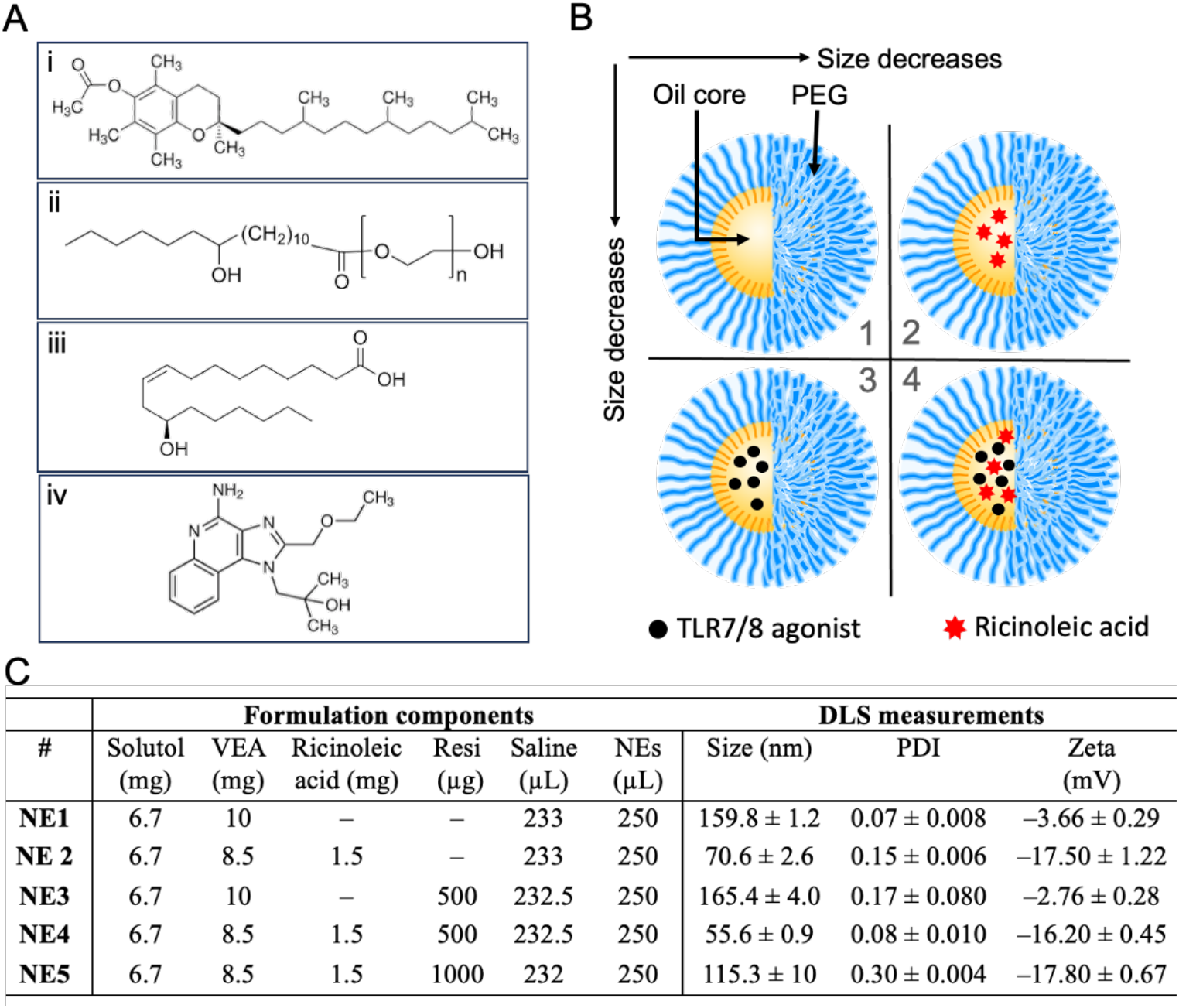
Nanoemulsion formulations. (a) Components of NEs; (i) vitamin E acetate (VEA), (ii) Solutol PEG surfactant, (iii) ricinoleic acid, and (iv) resiquimod drug. (b) Schematic representation of free NEs formulations consisting of an oil core and PEG solubilizer, along with their encapsulated ingredients NEs (1–4). (c) The table outlines NEs compositions, ratios, and DLS characterization, including size, PDI, and surface charge.

### Nanoemulsions Characterization

The physicochemical properties of the nanoemulsions were comprehensively evaluated using three complementary analytical techniques. Particle size distribution, polydispersity index (PDI), and zeta potential (*ζ*) were determined by dynamic light scattering (DLS) using a Zetasizer Nano-ZS (Malvern Instruments, UK). Measurements were performed at a standard concentration of 1 mg/mL, with triplicate readings to ensure reproducibility. For higher-resolution size analysis and particle concentration measurements, nanoparticle tracking analysis (NTA) was conducted using a ZetaView PMX 110 V3.0 system (Particle Metrix GmbH, Germany). Samples were prepared by 1:106 dilution in PBS followed by filtration through a 20 nm syringe filter, with measurements taken at 11 predetermined positions (two acquisitions each) under optimized instrument settings (sensitivity: 80; shutter: 100). Morphological characterization was achieved through transmission electron microscopy (TEM) using a LEO EM910 instrument (Carl Zeiss SMT Inc., Peabody, MA) operated at 80 kV. For TEM imaging, samples were diluted 100-fold, applied to copper/carbon grids, and negatively stained with 1% uranyl acetate to enhance contrast.

### Stability of R848-NEs in Fetal Bovine Serum

Nanoemulsion stability in fetal bovine serum (FBS) was assessed by incubating freshly prepared NEs in normal saline (0.9% NaCl) with 10% (v/v) FBS. Samples were maintained at 37 °C in a shaking incubator (200 rpm) for 48 h. Size distribution and PDI were monitored using DLS at multiple time points (0, 1, 2, 4, 6, 24, and 48 h).

### HPLC Analysis of R848 Drug

Resiquimod concentration was determined using reversed-phase high-performance liquid chromatography (HPLC) on an Agilent 1100 system equipped with a Nucleosil C18 column (5 µm, 250 mm × 4.6 mm, Supelco). Analysis was performed following previously established methods [29].

### Drug Loading Efficiency (LE) and Loading Capacity (LC)

Drug loading efficiency and capacity were calculated using the following equations:

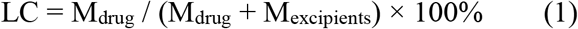

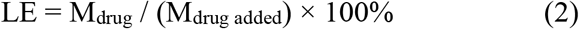

where M_drug_ represents the weight of the loaded drug, M_excipients_ refers to the excipient weight in the dispersion, and M_drug_ is the initially added drug weight.

### *In Vitro* Drug Release Studies

R848-loaded nanoemulsions (R848-NEs, 2 mg/mL) were subjected to release profiling under physiological sink conditions (PBS, 37 °C). Samples (100 µL) were loaded into pre-washed 3.5 kDa molecular weight cutoff Slide-A-Lyzer dialysis devices. Release kinetics were assessed at predetermined intervals (1, 2, 4, 6, and 24 h; n=3 replicates per timepoint). Following dialysis, samples were lyophilized, reconstituted in methanol (100 µL), and clarified by centrifugation. R848 quantification was performed via HPLC after salt precipitation, enabling precise determination of release kinetics.

### Cell Viability Studies

The cytotoxic effects of R848-loaded nanoemulsions (R848-NEs) were evaluated in both immune (RAW 264.7 macrophages) and tumor (4T1 TNBC) cell lines using complementary assays. For initial screening, cells were seeded in 96-well plates at 104 cells/well and allowed to adhere for 24 h. Serial dilutions of R848-NEs in complete medium were applied, and viability was quantified after 24 h using the CCK-8 assay, with data expressed as mean ± SD from six biological replicates.

To further validate dose-dependent effects, an MTT assay was performed on 4T1 cells (**Figure 7**). Cells were plated at 5,000 cells/well and cultured to 70–80% confluence. Treatments included R848-NEs (3.1– 100 µM), free resiquimod, or LPS (control) in complete medium (200 µL/well). After 24 h, cells were incubated with MTT solution (5 mg/mL, 20 µL/well) for 4 h at 37 °C. Formazan crystals were solubilized in DMSO (100 µL/well), and absorbance was measured at 590 nm (Bio-Rad microplate reader). Dose-response curves and IC_50_ values for NE3, NE4, and LPS were derived using CompuSyn software.

### Cellular Uptake of R848-NEs in RAW264.7 Macrophages

RAW 264.7 and NR-4T1 cells were plated at 105 cells/well in culture plates. For fluorescence tracking, NE3 and NE4 formulations (40 mg/mL) were labeled with DiI dye (4 µL of 25 mM solution) and purified via size-exclusion chromatography (Nab10 column) to yield 1.5 mL of dye-incorporated NEs. Cells were treated with 0.34 mg/mL DiI-NEs in DMEM for 4 or 24 h, followed by two warm DMEM washes. Nuclei were counterstained with Hoechst 33258 (2 µg/mL, 20 min), and cellular internalization was visualized by fluorescence microscopy.

### Experimental Design and Cell Line Treatment Protocol

4T1 cells were cultured in T-25 flasks until reaching 70-80% confluence, then divided into five groups (three flasks per group): Control (complete medium only), LPS-treated, R848-NE3 at IC_50_, R848-NE4 at IC_50_, and free resiquimod at IC_50_ [30, 31]. After 24 h of incubation, cells were harvested, and total protein content was measured using the Bradford assay [32]. Aliquots were stored at -80 °C for further analysis.

### Determination of Cell-Signaling Targets Using Enzyme-Linked Immunosorbent Assay (ELISA)

Protein expression levels of the cell signaling molecular targets and cell cycle regulators were detected in 4T1 cells exposed to different R848-NEs at IC_50_ doses of 24 h using ELISA technique. The detected proteins detected TLR4, TLR7, Cyclin D1, p65-NFκB, PD-L1 (CD274), (Casp-3), p62, LC3II, and Beclin-1 (BECN1).

## Results and discussion

### Preparation and characterization of Nes

We initially prepared nanoemulsions using a spontaneous emulsification method, as described in our previous works [25-28]. By employing a mix-and-match technique, we explored the solubility of various lipids and created a homogeneous oil phase. This was achieved through their physical and chemical interactions, including hydrophobic interactions, hydrogen bonding, and electrostatic forces. As a result, we obtained highly efficient, stable, long-lasting, reproducible, and scalable formulations. **Figure 1A** represents the chemical structures of components involved in the formulations NE1-4 (**Figure 1B**). A series of NE1-5 were prepared with and without TLR7/8 agonist at different concentrations and in the presence and absence of ricinoleic acid (**Figure 1C**). The data shows that ricinoleic acid decreased the particle sizes to nearly half, from 160 nm to 70 nm, likely due to physical and chemical interaction with other components. Similarly, it contracts the particle size and PDI of the nanodroplet emulsions encapsulated resiquimod drug as indicated in NE4 and NE5 for the same reason. Further, the presence of ricinoleic, a reactive lipophilic molecule, led to increasing the negative surface charge (zeta potential) due to the COOH groups presented at the oil/water interfacial surface.

The data presented in **Figure 2** highlight the robustness and consistency of the developed NE formulations. The DLS histograms of NE1–5 show narrowly distributed sizes and very low PDIs, indicating a uniform size distribution crucial for consistent behavior in biological systems. The low PDI confirms the homogeneity of the particle sizes, suggesting a highly controlled and reproducible formulation process. TEM images further reveal the spherical shape of the particles with high homogeneity and monodispersity, aligning with the DLS data and reinforcing the reliability of the findings. The consistency between the DLS and TEM results suggests that the morphological structure of NEs remains unaffected by the presence or absence of ricinoleic acid or resiquimod drug. This indicates that incorporating these components does not compromise the structural integrity of the nanoemulsions, which is essential for their stability and efficacy in drug delivery applications. Overall, the data demonstrate that the nanoemulsions maintain their desirable properties regardless of the inclusion of additional components, underscoring the robustness of the formulation.

**Figure 2.**
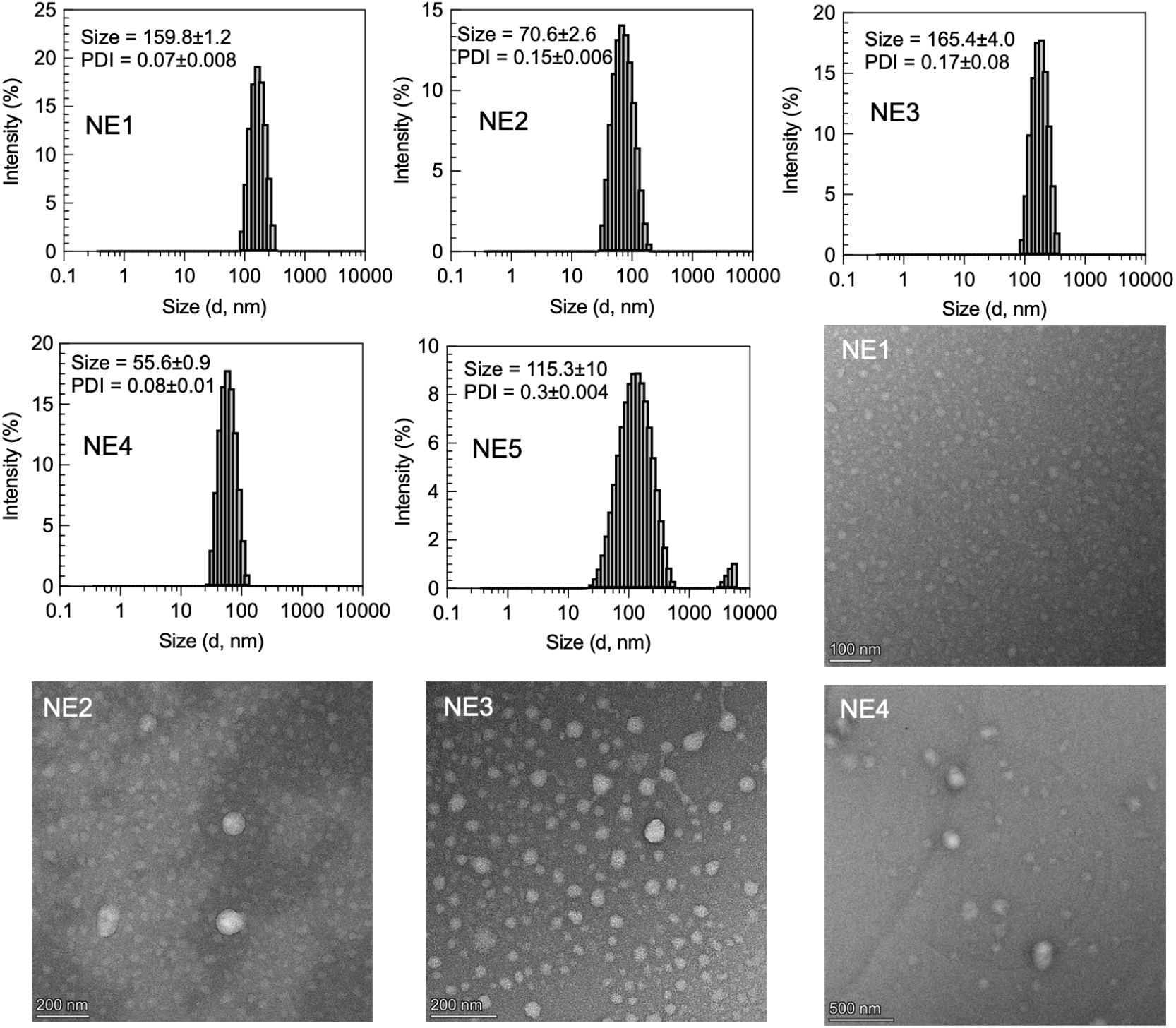
DLS histograms (NE1–5) and TEM morphology images for nanoemulsion formulations (NE1–4).

We evaluated the stability of the produced nanoemulsions in a biological environment using fetal bovine serum (FBS). NE1-5 were incubated in FBS for two days to examine their interaction with plasma proteins, driven by particle size agglomeration. The data showed remarkable size stability, with no changes observed in the size and PDI of all NEs, demonstrating significant monodispersity, integrity, and stability *in vitro* (**Figure 3A-B**). We also monitored the particle size by DLS over 20 days, storing the NEs at either room temperature or 4°C. The results showed negligible changes in particle size and PDI (**Figure 3C-F**), confirming high shelf-life stability and storage. This stability in both biological and storage conditions highlights the robustness of the NEs, making them promising candidates for drug delivery and other biomedical applications.

**Figure 3.**
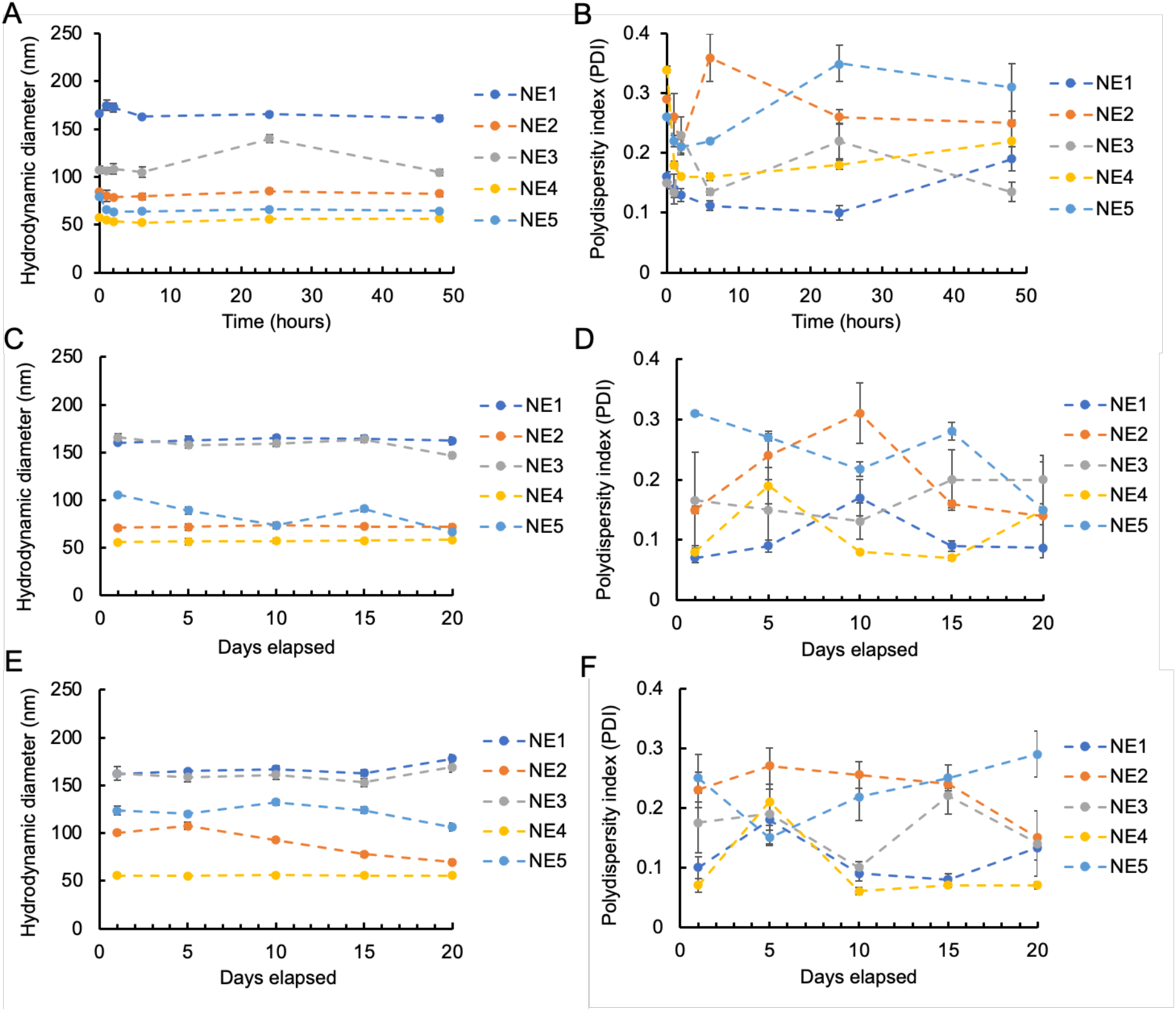
Stability of NE1-5. Size and PDI of NEs incubated in FBS over 48 hours at 37 °C (A-B). Size and PDI of NEs over time at RT (C-D) and 4 °C (E-F).

We validated the data using nanoparticle tracking analysis Zetaview to determine the size and particle concentration, which ranged from 3.7×10^13^ to 1.5×10^14^ particles/mL (**Figure 4A**). Next, we prepared and purified the nanoemulsions (NEs) to remove any drug that might be attached or adsorbed to the surface. The encapsulation efficiency (EE) and loading capacity (LC) of resiquimod drug within the nanoemulsions (NE3-5) were determined by HPLC, using equations 1 and 2 from the experimental section. The results showed EE of almost 100% and LC ranging from 2.9% to 5.6%, as shown in **Figure 4A**. We then investigated the drug release kinetics from NE3-5 using sink conditions in PBS at 37 °C for 24 h (**Figure 4B**). It was observed that the presence of ricinoleic acid in the formulations slowed the drug release, likely due to the possible interaction with the drug and other components of the nanoemulsions.

**Figure 4.**
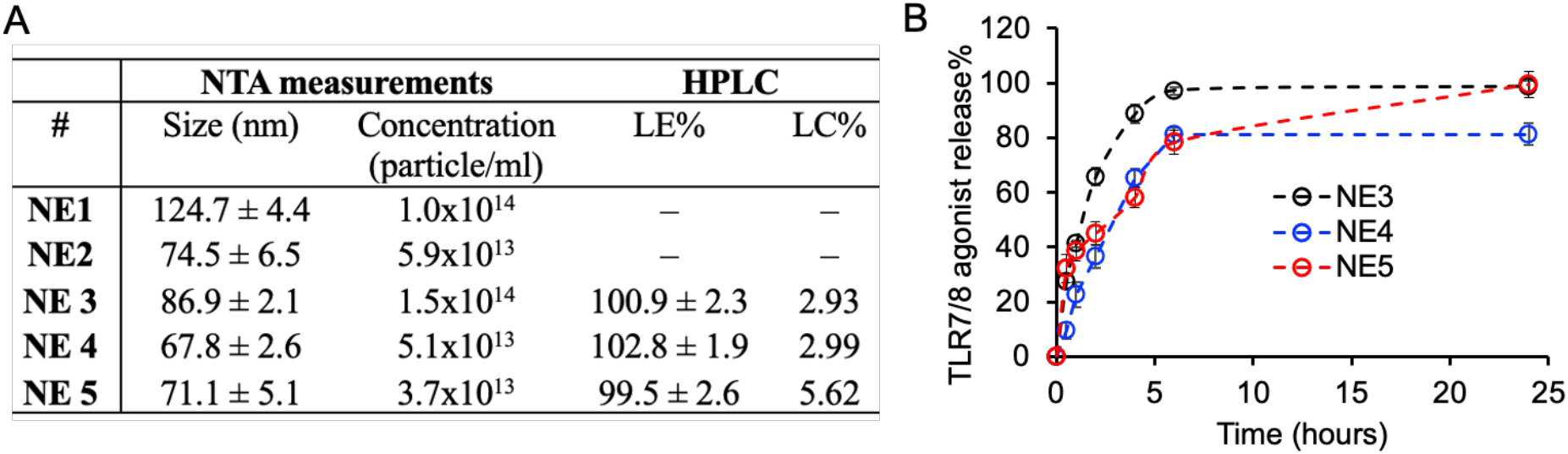
(A) NTA analysis, including size and particle concentration of NEs, along with HPLC to determine LE and LC of resiquimod. (B) drug release kinetics over 24 hours.

The cytotoxicity of all NE1-4 was evaluated using the CCK-8 assay in Raw 264.7 macrophages and 4T1 TNBC cell lines over a 24 h period to confirm their cytocompatibility *in vitro*. The data indicated that the presence of ricinoleic acid in both NE2 (0.68 mg NEs/mL, containing 20 µg resiquimod) and NE4 (1.36 mg NEs/mL, containing 40 µg resiquimod), resulted in significant cell death in both RAW macrophages and 4T1 cells (**Figure 5A**). Additionally, morphological changes were observed in RAW macrophages after treatment with either free NEs or Resi-NEs, potentially due to macrophage polarization from the M2 to M1 phenotype. In contrast, no morphological changes were detected in the 4T1 cell lines (**Figure 5B**). The differential response between RAW macrophages and 4T1 cells underscores the importance of considering cell type-specific effects when evaluating the biocompatibility of nanoemulsions.

**Figure 5.**
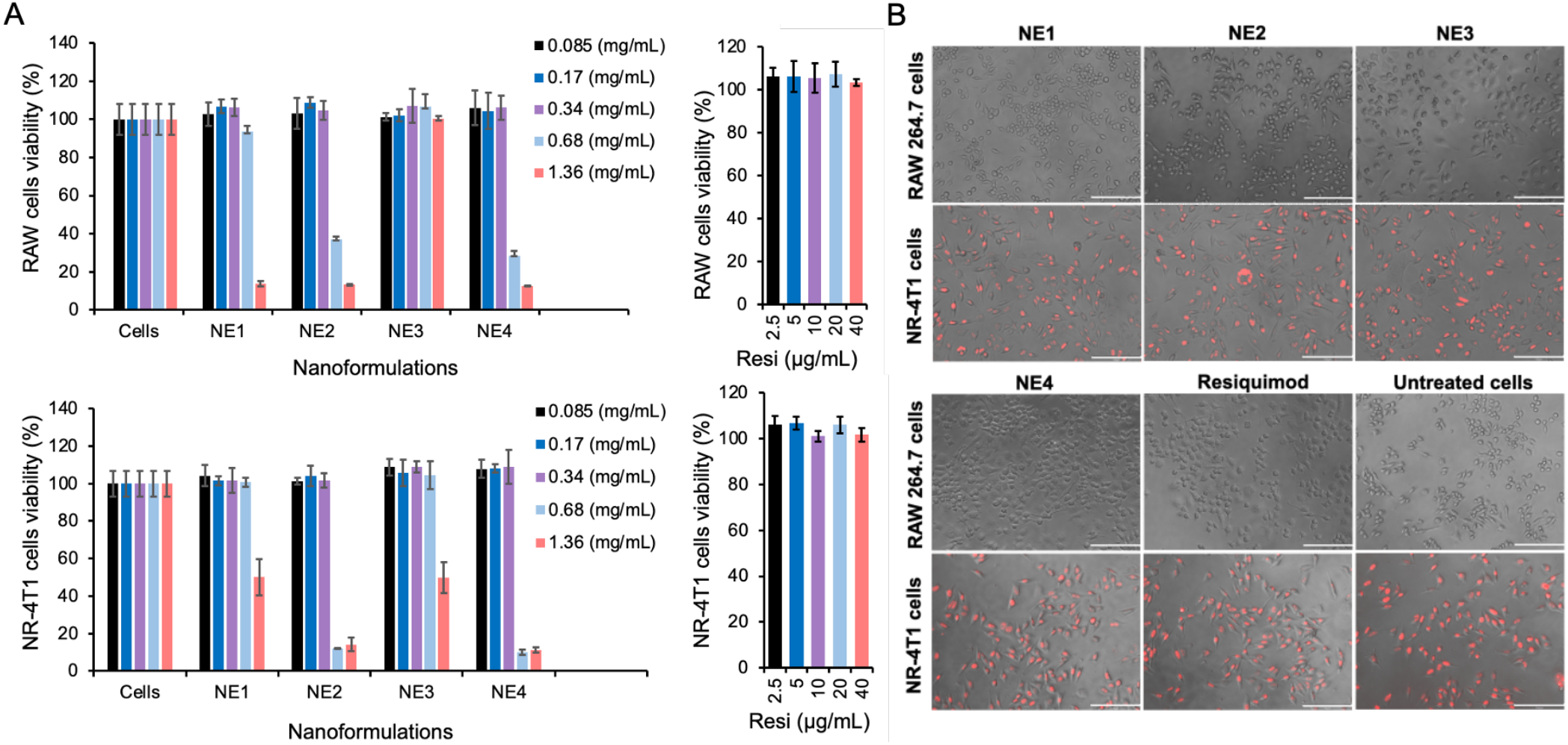
(A) CCK-8 assay for cytotoxicity measurements of NE1-4 in RAW 264.7 macrophages and NR-4T1 TNBC cancer cell lines. (B) Cell images were selected for a concentration of 0.34 mg/mL of NEs formulations. Scale bar is 150µm.

Cellular uptake experiments were performed on RAW macrophages and NR-4T1 cell lines by incubating fluorescently labeled NE3 and NE4 with the cells for 4 and 24 h. The uptake was visualized using fluorescence microscopy (**Figure 6A-C**). The images showed that both NEs were primarily localized in the cell membranes of RAW macrophages, with minimal entry into the cytoplasm at both time points (**Figure 6A and 6C(left)**). In contrast, the NEs were internalized into the cytoplasm of NR-4T1 cells (**Figure 6B and 6C(right)**). These results indicate that the NEs exhibit different uptake behaviors depending on the cell type. The significant localization in RAW macrophage membranes suggests a strong interaction with the cell surface, likely due to the phagocytic nature of macrophages. Conversely, the internalization into the cytoplasm of NR-4T1 cells suggests effective penetration and uptake by these cancer cells, which may have different endocytic pathways or a higher affinity for the NEs. These findings highlight the potential of NEs for targeted delivery in TNBC cells. Overall, the results confirm the safety profile of the formulated NEs *in vitro*, as they demonstrate appropriate cellular interactions without causing adverse effects.

**Figure 6.**
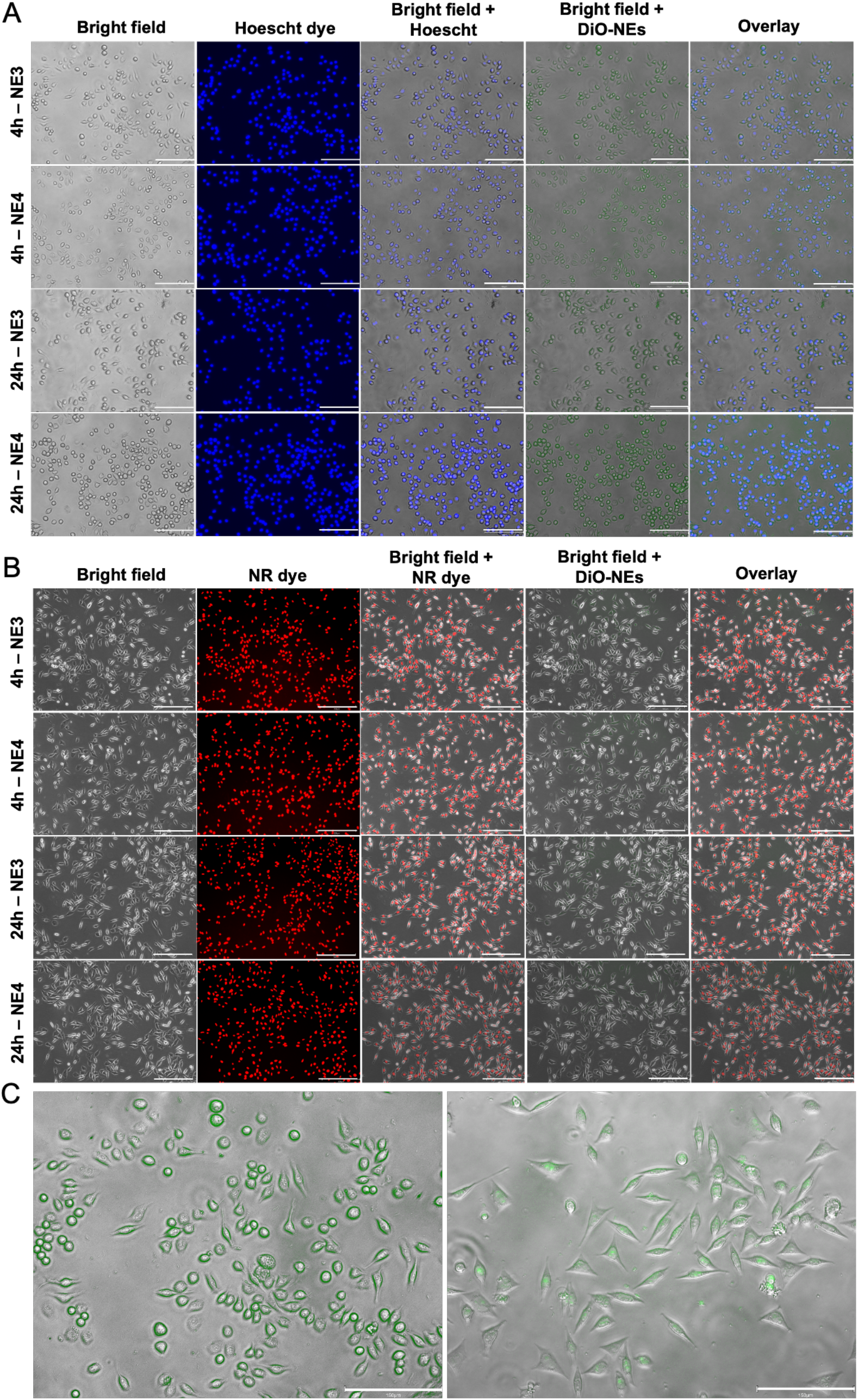
Cellular Uptake Data. Fluorescence images of RAW 264.7 macrophages (A) and NR-4T1 cells (B) showing the uptake of NE3 and NE4 at a concentration of 0.34 mg/mL at 4 and 24h incubation times. (C) Bright-field images of RAW macrophages (left) and 4T1 cells (right) demonstrating clear localization of LNE4 in the cellular membrane and cytoplasm, respectively. Scale bar is 150 µm.

We conducted an *in vitro* cytotoxicity assay to evaluate the growth inhibitory efficacy of different formulations of resiquimod and lipopolysaccharide (LPS) on 4T1 cells over 24 h, as shown in **Figure 7**. The IC_50_ of free resiquimod was 21.55 μM. However, they decreased to 14.95 μM and 11.34 μM of resiquimod in NE3 and NE4, respectively. The median growth inhibitory activity of LPS was 479.74 μM against 4T1 cells.

**Figure 7.**
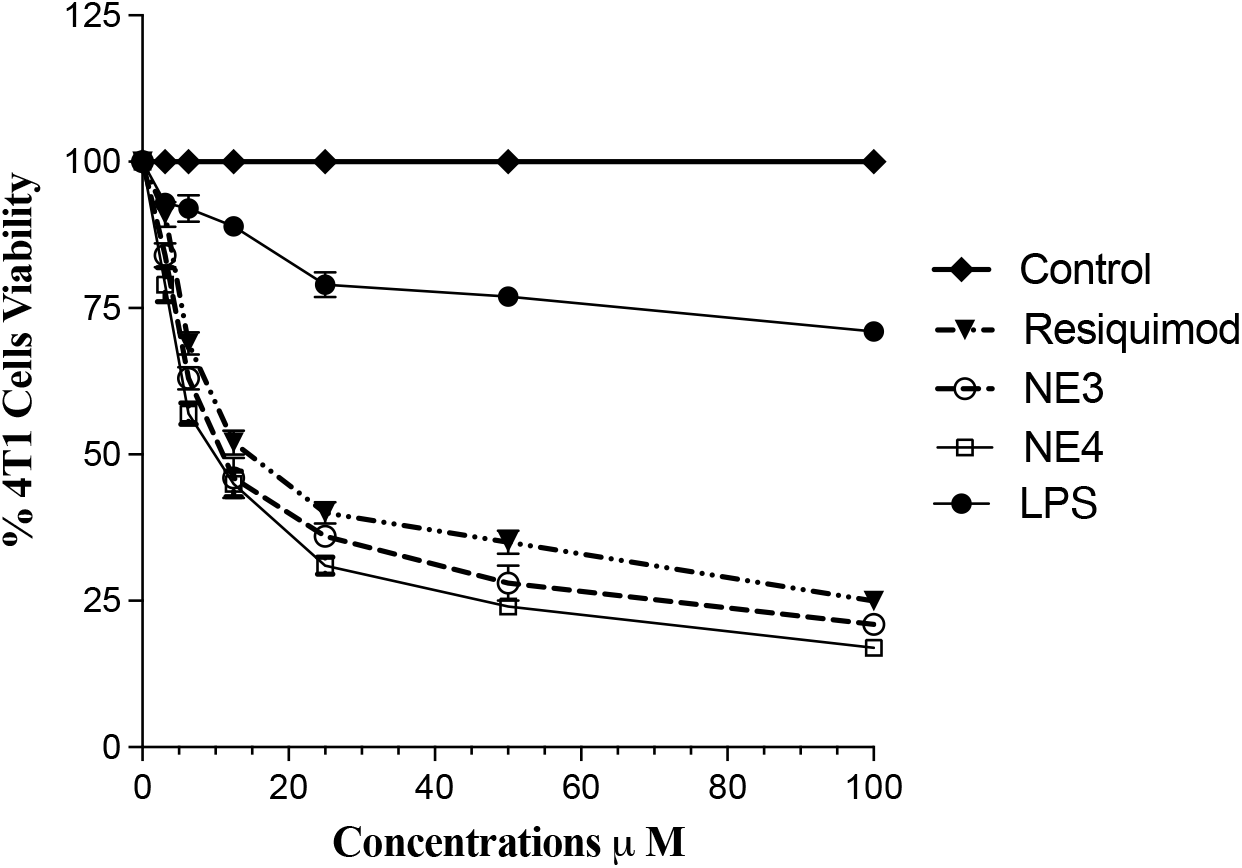
Cell viability using MTT assay of 4T1 cells with different treatments to determine the IC_50_ of resiquimod drug.

These findings revealed that resiquimod exerts a dose-dependent growth inhibitory effect on 4T1 cells, with a marked superiority of R848-NE4 nano-encapsulation. Similarly, evidence indicates that imiquimod (IMQ), another TLR7 agonist, inhibits the growth of various cancer types, including renal cell carcinoma [33]. intraepithelial neoplasia [34], squamous cell carcinoma [35], and basal cell carcinoma [36]. The anti-tumoral activities of TLR ligands have been demonstrated in various cancer cell types through indirect enhancement of the immune response activated by DCs and other innate and adaptive immune components within TME [37, 38]. However, the cellular mechanisms underlying TLR-induced anti-cancer activities remain unclear. Interestingly, some studies suggest a complex role for TLR7 in carcinogenesis and immunosuppression, warranting further investigation [39]. Our study aimed to investigate direct antitumor effects of different formulations of resiquimod, a TLR7 agonist, on 4T1 murine breast cancer cell line and explore the underlying molecular mechanisms.

We determined biochemical parameters in 4T1 cell line *in vitro*. **Figure 8** presents the levels of various cellular proteins and activities in different treatment groups: Control, resiquimod, NE3, NE4, and LPS. The proteins and activities measured include TLR7, TLR4, p65 NF-κB, Cyclin D1, Caspase-3 activity, PDL-1, Beclin 1, LC3 II, and p62. **Figure 8A-B** shows the levels of TLR7 and TLR4 expression in 4T1 control cells and those treated with different resiquimod formulations and LPS. No significant changes were observed in the basal expression of TLR4 across all treated 4T1 cells. In contrast, both LPS and resiquimod nano-encapsulation in NE3 or NE4 induced a slight increase in TLR7 protein expression. **Figure 8C** demonstrates that LPS, free resiquimod, and its nano-formulations NE3 and NE4, induce p65-NFκB activation compared to untreated cells. **Figure 8D-E** reports that neither LPS nor different formulations of resiquimod induced substantial changes in the proliferation marker Cyclin D1 or the apoptotic marker Caspase-3 compared to untreated cells. PD-L1 protein expression, acting as an immune checkpoint, was significantly increased in R848-NE4, showing a two-fold increase compared to untreated 4T1 cells and all other treatments, as shown in **Figure 8F**. The effects of different treatment regimens on autophagic markers Beclin-1, LC3II, and p62 are shown in **Figures G-I**, respectively. Significant induction of autophagy was observed in all resiquimod-treated 4T1 cells, whether free or in nano-formulations. However, the most marked induction of autophagy was observed in the R848-NE4 formula, evidenced by the significant increase in positive autophagic regulators Beclin-1 and LC3II and the marked inhibition of the negative autophagic regulator p62 compared to untreated 4T1 cells.

**Figure 8.**
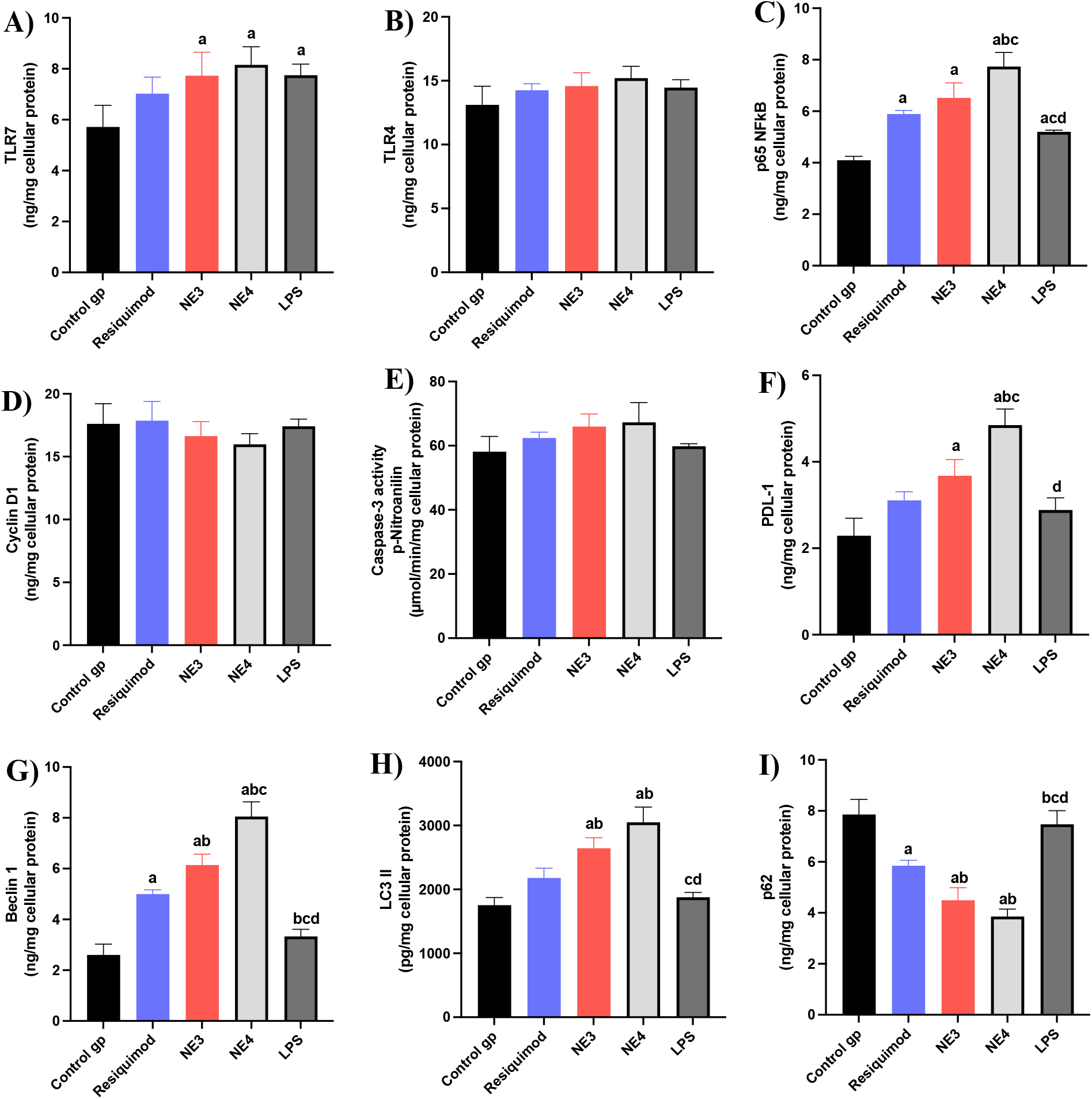
ELISA data was obtained from 4T1 breast cancer cells treated with NE formulations, compared to LPS-treated and untreated cells.

The findings indicate that resiquimod, through TLR7 stimulation, induces the expression of NF-κB and tumoral PD-L1 in the 4T1 breast cancer cell line. This induction is associated with the upregulation of autophagic markers, such as Beclin-1 and LC3II, and the inhibition of p62, without affecting cancer cell proliferation (as indicated by Cyclin D1 levels) or apoptosis (as indicated by Caspase-3 levels). The superiority of R848-NE4 was evident across all these molecular targets.

Growing evidence underscores the critical interplay between chronic inflammation, cell survival pathways, and cancer progression, with NF-κB serving as a pivotal regulatory hub [40]. While certain TLRs have demonstrated tumor-promoting effects through growth facilitation, the specific role of TLR7 remains underexplored [41]. Our findings suggest that resiquimod-mediated TLR7 activation triggers canonical NF-κB signaling via IκBα phosphorylation, followed by nuclear translocation of p50/p65 heterodimers and subsequent target gene transcription. These results corroborate earlier work by Cherfils-Vicini *et al*., wherein the TLR7 agonist loxoribine similarly activated NF-κB, upregulated the anti-apoptotic factor Bcl-2, and conferred both survival advantages and chemoresistance in human lung carcinoma models [42].

Resiquimod-induced autophagy may be partly attributed to the activated NF-κB signaling pathway in 4T1 cells. Consistent with our results, Cho *et al*. demonstrated that IMQ induces autophagy in mouse melanoma cells, associated with activated NF-κB signaling and overexpression of LC3II, Beclin-1, and other autophagic markers Atg 5-12 [43]. Furthermore, IMQ induced autophagic cell death in colon cells [44], and radioresistant MCF-7 breast cancer cells [45]. Interestingly, autophagy can have dual roles in cancer, acting as either a tumor suppressor or a survival mechanism depending on the context of the tumor microenvironment. Thus, our results raise the question of whether resiquimod-induced autophagy promotes cell survival rather than cancer cell death. However, the lack of impact on cancer cell proliferation and apoptosis suggests that resiquimod’s effects are primarily immunomodulatory rather than cytotoxic. Further investigations are highly recommended to uncover this crucial dogma.

The upregulation of PD-L1, a key immune checkpoint molecule, in resiquimod-treated 4T1 cells suggests that resiquimod may modulate the tumor microenvironment by enhancing immune evasion mechanisms. This effect could also be attributed to the activated NF-κB signaling pathway in 4T1 cells, consistent with findings reported by Zhang *et al*. [46]. The induction of PD-L1 expression by resiquimod indicates that coadministration with PD-L1 inhibitors may be a promising therapeutic strategy to enhance the antitumor immune response by preventing immune evasion. Further studies are needed to explore the synergistic effects of this combination therapy and determine the optimal dosing and scheduling for clinical application. In conclusion, our study highlights the potential of resiquimod as an immunostimulatory agent in cancer therapy and underscores the importance of combining it with PD-L1 inhibitors to achieve a more effective anti-tumor response.

## Conclusion

This study successfully developed and characterized two classes of R848-loaded nanoemulsions—neutral and ricinoleic acid-modified (negatively charged)—demonstrating that the incorporation of ricinoleic acid (particularly R848-NE4) significantly enhanced physicochemical stability, drug delivery efficiency, and biological performance compared to neutral formulations. Through systematic characterization (DLS, NTA, TEM) and in vitro evaluation, we established that R848-NE4 exhibits optimal particle properties, sustained release kinetics, and cell-type-specific uptake patterns—localizing to macrophage membranes while achieving cytoplasmic delivery in 4T1 tumor cells. Most notably, R848-NE4 triggered dual therapeutic effects: (1) autophagy-mediated tumor suppression (confirmed by LC3II/p62/Beclin-1 modulation) and (2) PD-L1 upregulation, suggesting a novel combinatorial strategy with immune checkpoint inhibitors. These findings position ricinoleic acid-stabilized nanoemulsions as a versatile platform for TNBC therapy, capable of simultaneously addressing tumor cell vulnerability (via autophagy induction) and immune evasion (through PD-L1 blockade). The demonstrated synergy between R848-NE4 and PD-1/PD-L1 axis modulation provides a strong rationale for further preclinical development and clinical translation of this approach.

## Acknowledgments

Mohamed Attia’s research is supported by the Carolina Cancer Nanotechnology Training Program, which is funded by NCI grant T32CA196589.

## References

[1] Y. Li et al., “Recent advances in therapeutic strategies for triple-negative breast cancer,” Journal of hematology & oncology, vol. 15, no. 1, p. 121, 2022.

[2] L. Yin, J.-J. Duan, X.-W. Bian, and S.-c. Yu, “Triple-negative breast cancer molecular subtyping and treatment progress,” Breast Cancer Research, vol. 22, pp. 1–13, 2020.

[3] A. Marra, D. Trapani, G. Viale, C. Criscitiello, and G. Curigliano, “Practical classification of triple-negative breast cancer: intratumoral heterogeneity, mechanisms of drug resistance, and novel therapies,” NPJ breast cancer, vol. 6, no. 1, p. 54, 2020.

[4] Z. Guo et al., “Tumor microenvironment and immunotherapy for triple-negative breast cancer,” Biomarker Research, vol. 12, no. 1, pp. 1–19, 2024.

[5] G. K. Gupta et al., “Perspectives on triple-negative breast cancer: current treatment strategies, unmet needs, and potential targets for future therapies,” Cancers, vol. 12, no. 9, p. 2392, 2020.

[6] M. Nedeljković and A. Damjanović, “Mechanisms of chemotherapy resistance in triple-negative breast cancer—how we can rise to the challenge,” Cells, vol. 8, no. 9, p. 957, 2019.

[7] J. Zhou, L. Zhang, S. Liu, D. DeRubeis, and D. Zhang, “Toll-like receptors in breast cancer immunity and immunotherapy,” Frontiers in Immunology, vol. 15, p. 1418025, 2024.

[8] Y. Abdou, A. Goudarzi, J. Yu, S. Upadhaya, B. Vincent, and L. Carey, “Immunotherapy in triple negative breast cancer: beyond checkpoint inhibitors. Npj Breast Cancer. 2022; 8: 121,” ed: Doi.

[9] H. Sun et al., “Targeting toll-like receptor 7/8 for immunotherapy: recent advances and prospectives,” Biomarker research, vol. 10, no. 1, p. 89, 2022.

[10] S. Chakraborty et al., “Application of toll-like receptors (TLRs) and their agonists in cancer vaccines and immunotherapy,” Frontiers in Immunology, vol. 14, p. 1227833, 2023.

[11] K. R. Bhaliya, M. Anwer, A. Munn, and M. Q. Wei, “New horizons in cancer immunotherapy: The evolving role of R848 and R837,” Molecular and Clinical Oncology, vol. 22, no. 1, p. 4, 2024.

[12] J. Ye et al., “Toll-like receptor 7/8 agonist R848 alters the immune tumor microenvironment and enhances SBRT-induced antitumor efficacy in murine models of pancreatic cancer,” Journal for Immunotherapy of cancer, vol. 10, no. 7, p. e004784, 2022.

[13] J. Zhou, Y. Xu, G. Wang, T. Mei, H. Yang, and Y. Liu, “The TLR7/8 agonist R848 optimizes host and tumor immunity to improve therapeutic efficacy in murine lung cancer,” International Journal of Oncology, vol. 61, no. 1, p. 81, 2022.

[14] K. Pfleiderer et al., “NIMG-48. TLR7/8-AGONIST-LOADED NANOPARTICLES REPROGRAM TUMOR-ASSOCIATED MYELOID CELLS FOR EFFECTIVE IMMUNOTHERAPY OF EXPERIMENTAL GLIOMA AND MRI-BASED TREATMENT MONITORING,” Neuro-Oncology, vol. 23, no. Supplement_6, pp. vi139–vi140, 2021.

[15] C. Hotz et al., “Reprogramming of TLR7 signaling enhances antitumor NK and cytotoxic T cell responses,” Oncoimmunology, vol. 5, no. 11, p. e1232219, 2016.

[16] H. Chi et al., “Anti-tumor activity of toll-like receptor 7 agonists,” Frontiers in pharmacology, vol. 8, p. 304, 2017.

[17] M. Luchner, S. Reinke, and A. Milicic, “TLR agonists as vaccine adjuvants targeting cancer and infectious diseases. Pharmaceutics. 2021; 13 (2): 142, ” PUBMED.

[18] T. Zhao et al., “Vaccine adjuvants: mechanisms and platforms,” Signal transduction and targeted therapy, vol. 8, no. 1, p. 283, 2023.

[19] X. Su et al., “Strategies to enhance the therapeutic efficacy of anti-PD-1 antibody, anti-PD-L1 antibody and anti-CTLA-4 antibody in cancer therapy,” Journal of Translational Medicine, vol. 22, no. 1, p. 751, 2024.

[20] S. A. Weiss, J. D. Wolchok, and M. Sznol, “Immunotherapy of melanoma: facts and hopes,” Clinical Cancer Research, vol. 25, no. 17, pp. 5191–5201, 2019.

[21] A. Knight, L. Karapetyan, and J. M. Kirkwood, “Immunotherapy in melanoma: recent advances and future directions,” Cancers, vol. 15, no. 4, p. 1106, 2023.

[22] S. Bhagchandani, J. A. Johnson, and D. J. Irvine, “Evolution of Toll-like receptor 7/8 agonist therapeutics and their delivery approaches: From antiviral formulations to vaccine adjuvants,” Advanced drug delivery reviews, vol. 175, p. 113803, 2021.

[23] G. Frega et al., “Trial Watch: experimental TLR7/TLR8 agonists for oncological indications,” Oncoimmunology, vol. 9, no. 1, p. 1796002, 2020.

[24] C. Rolfo, E. Giovannetti, P. Martinez, S. McCue, and A. Naing, “Applications and clinical trial landscape using Toll-like receptor agonists to reduce the toll of cancer,” npj Precision Oncology, vol. 7, no. 1, p. 26, 2023.

[25] M. F. Attia, M. I. Swasy, R. Akasov, F. Alexis, and D. C. Whitehead, “Strategies for High Grafting Efficiency of Functional Ligands to Lipid Nanoemulsions for RGD-Mediated Targeting of Tumor Cells In Vitro,” ACS Applied Bio Materials, vol. 3, no. 8, pp. 5067–5079, 2020.

[26] M. F. Attia et al., “Radiopaque Iodosilane-Coated Lipid Hybrid Nanoparticle Contrast Agent for Dual-Modality Ultrasound and X-ray Bioimaging,” ACS Applied Materials & Interfaces, vol. 14, no. 49, pp. 54389–54400, 2022.

[27] M. F. Attia et al., “Functionalizing nanoemulsions with carboxylates: impact on the biodistribution and pharmacokinetics in mice,” Macromolecular Bioscience, vol. 17, no. 7, p. 1600471, 2017.

[28] M. F. Attia et al., “Functionalization of nano-emulsions with an amino-silica shell at the oil– water interface,” RSC advances, vol. 5, no. 91, pp. 74353–74361, 2015.

[29] N. Vinod et al., “High-capacity poly (2-oxazoline) formulation of TLR 7/8 agonist extends survival in a chemo-insensitive, metastatic model of lung adenocarcinoma,” Science advances, vol. 6, no. 25, p. eaba5542, 2020.

[30] F. M. Abdallah, A. I. Ghoneim, M. M. Abd-Alhaseeb, I. T. Abdel-Raheem, and M. W. Helmy, “Unveiling the antitumor synergy between pazopanib and metformin on lung cancer through suppressing p-Akt/NF-κB/STAT3/PD-L1 signal pathway,” Biomedicine & Pharmacotherapy, vol. 180, p. 117468, 2024.

[31] S. M. El-Hanboshy, M. W. Helmy, and M. M. Abd-Alhaseeb, “Catalpol synergistically potentiates the anti-tumour effects of regorafenib against hepatocellular carcinoma via dual inhibition of PI3K/Akt/mTOR/NF-κB and VEGF/VEGFR2 signaling pathways,” Molecular biology reports, vol. 48, pp. 7233–7242, 2021.

[32] F. He, “Bradford protein assay,” Bio-protocol, pp. e45–e45, 2011.

[33] E. C. Kauffman, H. Liu, M. J. Schwartz, and D. S. Scherr, “Toll-Like Receptor 7 Agonist Therapy with Imidazoquinoline Enhances Cancer Cell Death and Increases Lymphocytic Infiltration and Proinflammatory Cytokine Production in Established Tumors of a Renal Cell Carcinoma Mouse Model,” Journal of oncology, vol. 2012, no. 1, p. 103298, 2012.

[34] M. I. van Poelgeest et al., “Detection of human papillomavirus (HPV) 16-specific CD4+ T-cell immunity in patients with persistent HPV16-induced vulvar intraepithelial neoplasia in relation to clinical impact of imiquimod treatment,” Clinical cancer research, vol. 11, no. 14, pp. 5273–5280, 2005.

[35] M. Y. Ahn et al., “Toll-like receptor 7 agonist, Imiquimod, inhibits oral squamous carcinoma cells through apoptosis and necrosis,” Journal of oral pathology & medicine, vol. 41, no. 7, pp. 540–546, 2012.

[36] R. L. Miller, T.-C. Meng, and M. A. Tomai, “The antiviral activity of Toll-like receptor 7 and 7/8 agonists,” Drug news & perspectives, vol. 21, no. 2, pp. 69–87, 2008.

[37] B. Huang, J. Zhao, J. Unkeless, Z. Feng, and H. Xiong, “TLR signaling by tumor and immune cells: a double-edged sword,” Oncogene, vol. 27, no. 2, pp. 218–224, 2008.

[38] M. Damo, D. S. Wilson, E. Simeoni, and J. A. Hubbell, “TLR-3 stimulation improves anti-tumor immunity elicited by dendritic cell exosome-based vaccines in a murine model of melanoma,” Scientific reports, vol. 5, no. 1, p. 17622, 2015.

[39] T. Duan, Y. Du, C. Xing, H. Y. Wang, and R.-F. Wang, “Toll-like receptor signaling and its role in cell-mediated immunity,” Frontiers in Immunology, vol. 13, p. 812774, 2022.

[40] M. H. Park and J. T. Hong, “Roles of NF-κB in cancer and inflammatory diseases and their therapeutic approaches,” Cells, vol. 5, no. 2, p. 15, 2016.

[41] X. Chen, Y. Zhang, and Y. Fu, “The critical role of Toll-like receptor-mediated signaling in cancer immunotherapy,” Medicine in Drug Discovery, vol. 14, p. 100122, 2022.

[42] J. Cherfils-Vicini et al., “Triggering of TLR7 and TLR8 expressed by human lung cancer cells induces cell survival and chemoresistance,” The Journal of clinical investigation, vol. 120, no. 4, pp. 1285–1297, 2010.

[43] J. H. Cho et al., “The TLR7 agonist imiquimod induces anti-cancer effects via autophagic cell death and enhances anti-tumoral and systemic immunity during radiotherapy for melanoma,” Oncotarget, vol. 8, no. 15, p. 24932, 2017.

[44] J. Y. Yi, Y.-J. Jung, S. S. Choi, J. Hwang, and E. Chung, “Autophagy-mediated anti-tumoral activity of imiquimod in Caco-2 cells,” Biochemical and biophysical research communications, vol. 386, no. 3, pp. 455–458, 2009.

[45] S.-J. Kang, J.-H. Tak, J.-H. Cho, H.-J. Lee, and Y.-J. Jung, “Stimulation of the endosomal TLR pathway enhances autophagy-induced cell death in radiotherapy of breast cancer,” Genes & Genomics, vol. 32, pp. 599–606, 2010.

[46] H. Zhang et al., “Regulatory mechanisms of immune checkpoints PD-L1 and CTLA-4 in cancer,” Journal of Experimental & Clinical Cancer Research, vol. 40, no. 1, p. 184, 2021.

